# Don’t Stress, It’s Under Control: Neural Correlates of Stressor Controllability in Humans

**DOI:** 10.1101/2021.03.30.437657

**Authors:** Laura E. Meine, Jana Meier, Benjamin Meyer, Michèle Wessa

## Abstract

Animal research has repeatedly shown that experience of control over an aversive event can protect against the negative consequences of later uncontrollable stress. Neurobiologically, this effect is assumed to correspond to persistent changes in the pathway linking the ventromedial prefrontal cortex (vmPFC) and the dorsal raphe nucleus. However, it remains unclear to what extent these findings translate to humans. During functional magnetic resonance imaging, we subjected participants to controllable and uncontrollable aversive but non-painful electric stimuli, as well as to a control condition without aversive stimulation. In each trial, a symbol signalled whether participants could terminate the stressor through correct performance in a button-matching task or whether the stressor would be randomly terminated, i.e., uncontrollable. Along with neural responses, we assessed participants’ accuracy, reaction times, and heart rate. To relate neural activations and subjective experience, we asked participants to rate perceived control, helplessness, and stress. Results were largely in line with our hypotheses. The vmPFC was generally deactivated by stress, but this effect was attenuated when participants could terminate the stressor compared to when their responses had no effect. Furthermore, activation in stress-responsive regions, including the bilateral insula, was reduced during controllable trials. Under uncontrollable stress, greater vmPFC recruitment was linked to reduced feelings of helplessness. An investigation of condition-dependent differences in vmPFC connectivity yielded no significant results. Our findings further corroborate animal research and emphasise the role of the vmPFC in controllability-dependent regulation of stress responses. Based on the results, we discuss future directions in the context of resilience research and mental health promotion.

## 1. Introduction

The reaction to and appraisal of an aversive event is closely linked to actual or perceived control over the event (Steptoe & Poole, 2016). Specifically, animal studies have compellingly demonstrated that uncontrollable stress induces a failure to escape aversive stimulation and increases anxiety (Overmier & Seligman, 1967; Chourbaji et al, 2005; Maier & Seligman, 2016). This finding has been formalised in the prominent *theory of learned helplessness* which posits that a passive response to an inescapable stressor may generalise and thus render the individual helpless in the face of subsequent aversive stimulation, even if they could escape it (Overmier & Seligman, 1967; Maier & Seligman, 2016). In contrast, the experience of control over a stressor was shown to be protective. Animals that could terminate aversive stimulation were not susceptible to subsequent uncontrollable stress, as evidenced by e.g., lack of freezing (Amat et al., 2010; Maier & Seligman, 2016). To harness the potential of this latter effect, its underlying neurobiological mechanisms have already been delineated in animals (Amat et al., 2005, 2006, 2008; Baratta et al., 2008; Maier & Watkins, 2010, Maier & Seligman, 2016). The results suggest that sensitisation of a pathway connecting the dorsal raphe nucleus (DRN) with the amygdala might explain the symptoms of learned helplessness. Building on these findings, the researchers targeted the ventromedial prefrontal cortex (vmPFC) as a cortical input structure to the DRN. They describe how the potential for control registers in the vmPFC which then inhibits the DRN via glutamatergic projections to GABAergic interneurons. In turn, serotonergic signalling from the DRN to the amygdala (and other stress-related regions) is reduced. Because changes in the vmPFC-DRN pathway persist, the individual appears *immunised* against later uncontrollable stress.

Since the first findings on differential effects of stressor controllability in animals emerged, researchers have investigated translations to humans, placing particular focus on learned helplessness (Thornton & Jacobs, 1971; Abramson et al., 1978; Taylor et al., 2014; Meine et al., 2020). Consequently, the effect of uncontrollable stress has long been discussed in terms of depression pathogenesis (Pryce et al., 2011). More recently, Henderson et al. (2012) could show a beneficial effect of control over a noise stressor on executive control functions. Similarly, Hartley et al. (2014) noted improvements in fear extinction following the experience of control over a stressor. Despite this, only few studies have addressed stressor controllability effects on a neural level in humans. A direct test of the model put forward by Maier and Seligman (2016), using neuroimaging techniques, proves difficult due to the small size of the DRN (Kranz et al., 2012). Existing studies therefore tend to focus on the vmPFC and stress-related brain regions. In the context of pain processing, Salomons et al. (2015) described increased vmPFC-amygdala connectivity under controllable pain and Bräscher et al. (2016) reported a pain-inhibiting function of the dorsolateral prefrontal cortex for controllable heat stimuli. Kerr et al. (2012) examined participants’ response to aversive videos which could either be avoided or not. They showed that vmPFC activity increases under anticipation of control and exerts an inhibitory effect on the amygdala. In line with this, Cremers et al. (2020) reported greater vmPFC efficiency in participants who could avoid mild shocks compared to another group who could not. A very recent study showed that vmPFC activation was predictive of recovered active avoidance behaviour following passivity under uncontrollable stress (Wang & Delgado, 2021). Others observed diminished stress effects on the vmPFC only if threats were both controllable and predictable (Wood et al., 2015). Using a fear conditioning paradigm, Wanke & Schwabe (2020b) directly contrasted instrumental control with passive extinction and found more pronounced fear reduction in the former condition. However, they noted increased activation of the vmPFC under uncontrollable stress, but not under instrumental control. Another recent study showed the expected decrease in activation of threat-responsive brain regions when participants could terminate mild electric shocks, however, they observed no involvement of the vmPFC (Limbachia et al., 2021). Inconsistencies in results may at least in part be explained by differences in experimental designs (e.g., between- vs. within-subject) and the type of stressor used (e.g., electric shocks, thermal stimulation, noise, or social-evaluative stress). Many have studied aversive stimuli which are signalled by a cue and can be avoided (Diener et al., 2009; Kerr et al., 2012; Wanke & Schwabe, 2020b; Cremers et al., 2020). However, we feel that researchers should also investigate controllability-dependent responses in human behaviour and brain function under acute stress. Since stress describes an omnipresent fact of life, and its avoidance may not be possible under all circumstances, the study of factors or mechanisms that can alleviate stress effects certainly presents an ecologically valid research endeavour.

In the present study, we therefore aimed at further elucidating the neural correlates of control over acute stress in humans. We chose a within-subject design to maximise comparability of conditions and focus on immediate effects of controllability, as opposed to effects on post-stress read-outs. To this end, we conducted an event-related functional magnetic resonance imaging (fMRI) experiment in which participants were subjected to controllable stress, uncontrollable stress, and a baseline condition without aversive stimulation. Along with neural activation, performance and reaction times (RT) were assessed and heart rate was recorded. To allow associations between neurophysiological results and participants’ subjective experience, they also supplied ratings on perceived control, helplessness, and stress. We expected to observe effects of both stress and controllability on outcomes of interest. Specifically, we hypothesised that, compared to baseline, participants would react more slowly, make more errors, report higher stress ratings, show enhanced activation of stress-related brain regions (e.g., amygdala, insula), and elevated heart rate under stress (controllable and uncontrollable). Concerning stressor controllability, we expected faster and more accurate responses, higher ratings of perceived control, but lower stress and helplessness ratings, greater vmPFC activation along with reduced activation in stress-activated regions, and lower heart rate in controllable compared to uncontrollable experimental trials. As DRN imaging in humans remains a challenge (Kranz et al., 2012), we only anticipated to observe controllability-dependent differences in vmPFC-amygdala connectivity.

## 2. Materials and methods

### 2.1 Participants

52 participants aged 19-30 took part in this study. Prior to data collection, all were screened for acute or chronic physical diseases as well as current or past Diagnostic and Statistical Manual of Mental Disorders (DSM)-IV axis I disorders using a semi-structured telephone interview similar to the Structured Clinical Interview for DSM-IV axis I disorders (SCID-I). We included only healthy participants who were fluent in German, MR compatible, right-handed, reported a body mass index (kg/m2) between 18.5 and 26, had no history of mental disorders, and took no psychopharmaceuticals. To prevent adverse health effects of our stress induction procedure, we confirmed that participants were neither pregnant, nor suffering from critical cardiac problems, or diagnosed with migraine. Seven participants were excluded from analysis: one broke off testing, two figured out the experimental manipulation (see manipulation check in the results section for details), one reported stimulus electrode malfunctioning, and three rated stressor aversiveness very low (< 25/100 in more than half the runs), also suggesting electrode malfunction/loosening. The remaining sample consisted of 45 participants (47% female, age: *M* = 24.64, *SD* = 3.05). For fMRI analyses, another participant was disregarded due to excessive motion (> 2mm translation, > 2° rotation between volumes) and a further two were excluded because of data storage problems (missing data).

This study was approved by the ethics committee of the Institute of Psychology, Johannes Gutenberg-University, Mainz, Germany (2017-JGU-psychEK-003, 26/5/2017), and was conducted according to the Declaration of Helsinki.

### 2.2 Procedure

Participants attended a 2 h MRI session. Upon arrival, they received information on the study along with a brief overview of procedures and provided written informed consent. Participants then completed a questionnaire on stress experiences and current well-being and entered the scanner. Inside the scanner, electric stimuli were calibrated, and participants underwent the stress induction procedure. During the stress induction, their heart rate (i.e., beats per minute; BPM) was tracked at a sampling rate of 50 Hz using a MR-compatible pulse-oximeter with an infrared emitter fixed to the participant’s left index finger. Participants were remunerated with 10€/hour or – if preferred – received course credit.

### 2.3 Stress induction

Stress was induced through individually calibrated intermittent double electric stimuli (two stimuli 20 ms apart every 1000 ± 250 ms for the duration of the stress phase). These were administered via Digitimer DS7A current stimulator through an electrode attached to the participant’s right ankle. After determining the perception threshold, stimulus intensity was increased in 0.5 mA increments. Participants rated each stimulus on a scale with four anchors: “barely perceptible”, “unpleasant”, “very unpleasant”, and “really painful”. We emphasised natural diversity in stimulus perception and encouraged honest responses. Calibration concluded with the participant reaching a very unpleasant, yet not painful intensity (5-6 out of 10; *M* = 5.33, *SD* = 0.58). This level was maintained for the duration of the experiment. To avoid startle responses and movement, participants were familiarised with the intermittent pattern of the electric stimuli. In contrast to distinct single stimuli, the relatively high-frequency stimulation represented a more continuous stress phase.

In an event-related fMRI design, adapted from a behavioural experiment (Meine et al., 2020), participants underwent four runs, each comprising 12 controllable stress (CON) trials, 12 uncontrollable stress (UNCON) trials, and 6 baseline trials without aversive stimulation (Figure 1). For each run, we created a pseudorandomised trial sequence using optseq2 (Dale, 1999; http://surfer.nmr.mgh.harvard.edu/optseq/) to consider slow changes in hemodynamic responses. Optseq2 effectively decorrelates regressors by shifting the inter-trial interval (ITI). In each trial, participants first saw a symbol indicating the trial type (! for CON, ? for UNCON, and a dot for baseline trials; 1500 ms) before a fixation cross appeared and aversive stimulation set in. Participants received intermittent electric stimuli for 4000 ms before a cue (circle, square, or triangle; distributed equally within condition and run) prompted them to press a corresponding button (index, middle, or ring finger of the right hand) to terminate the stimulation. Button presses were registered on a Current Designs response pad with four buttons of which participants were instructed to use only three. A feedback phase (3000 ms) followed, then a variable ITI (1000-6000 ms; *M* = 2000 ms, *SD* = 1280). In CON trials, correct responses given within 1000 ms immediately terminated the aversive stimuli with a happy smiley appearing on-screen. Responses that were incorrect or too slow led to continued stimulation throughout the feedback phase in which a sad smiley was displayed. In UNCON trials, positive feedback (i.e., termination of electric stimuli) was randomly given in 50% of trials, irrespective of the participant’s response. Baseline trials corresponded to CON trials excluding the aversive stimuli. Before the first run, participants completed a 12-trial stress-free training phase in which they learned the correct cue-button assignment (counterbalanced across participants), showing at least 80% correct performance. They were told that, in CON trials, stressor termination directly depended upon their response, whereas, in UNCON trials, it was up to chance. Nonetheless, we explicitly instructed participants to respond as quickly and correctly as possible in all trials.

**Figure 1.**
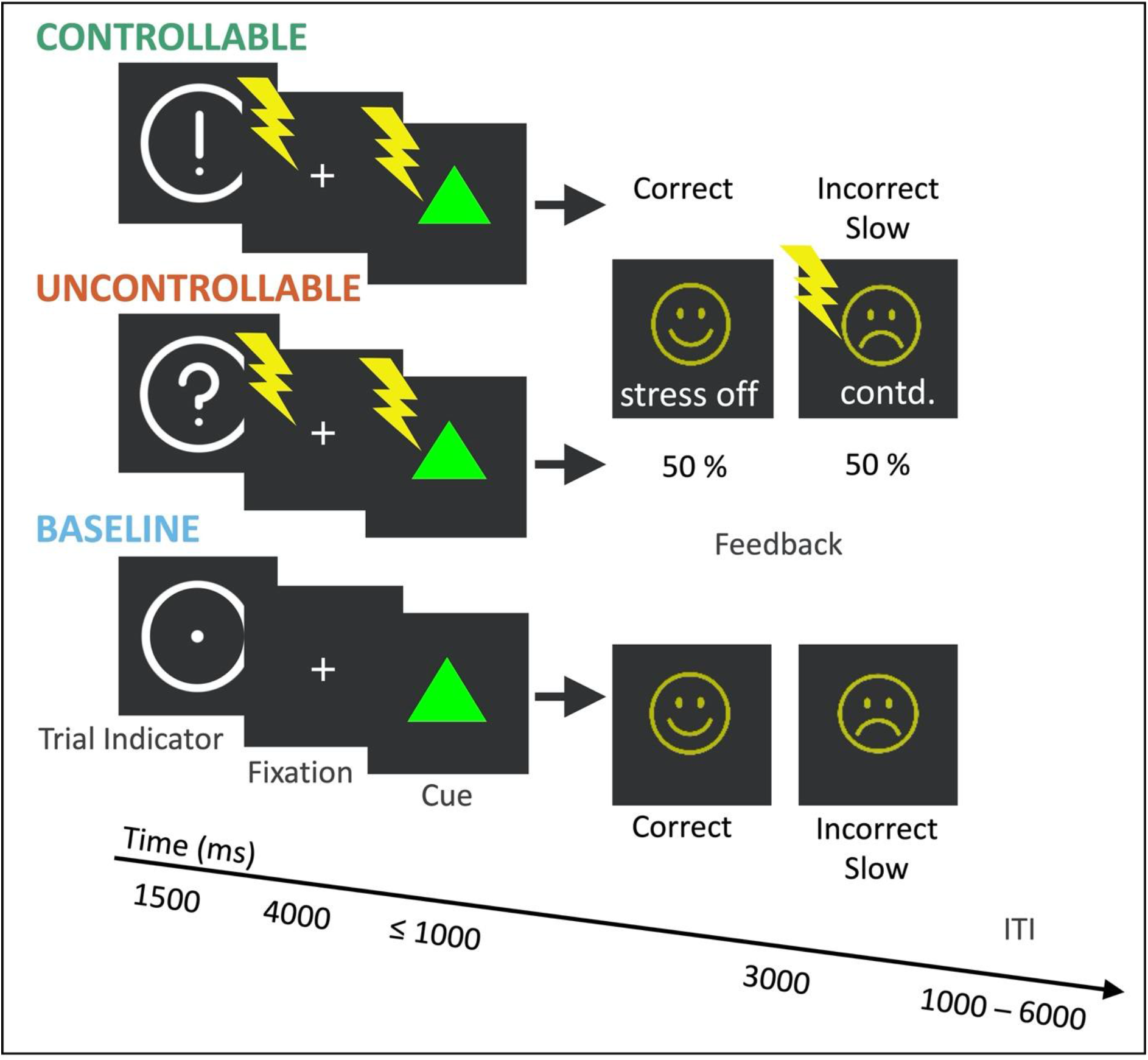
Stress induction paradigm: in controllable stress trials, participants’ response to a cue was instrumental for stressor termination. In uncontrollable stress trials, aversive stimulation was terminated randomly in 50% of trials, irrespective of participants’ response. Trials without stress served as baseline.

At the end of each run, participants indicated how stressed they had felt in each condition and – only for CON and UNCON – how much control over the stressor they had perceived and how helpless they had felt. Additionally, following the eighth CON and UNCON trial of each run, they rated stressor aversiveness. A scale from 0 (not at all) to 100 (very) in increments of 5 was used.

Given our manipulation, participants should have received on average more electric stimuli in UNCON (50% negative feedback) compared to CON trials (potentially 0% negative feedback). However, to ensure equal exposure, we extended the stress-phase prior to the cue in CON trials using a trial-by-trial approach. Specifically, correct responses in CON trials were tracked and the corresponding time required to match negative feedback in UNCON trials (6 x 3000 s in each run) was divided up and added to subsequent CON trials in a systematic fashion so as to make it barely noticeable. Other studies have followed a similar approach (Diener et al., 2009). Immediately after they had left the scanner, we asked participants if they had noticed any differences in the total number of electric stimuli between conditions.

This experiment was programmed in Python 2.7 (Van Rossum & Drake, 2010), using mainly the PsychoPy package (Peirce et al., 2019).

### 2.4 FMRI data acquisition and preprocessing

Structural and fMRI data were acquired using a 3T Siemens Trio Scanner with a 32-channel head coil. First, T2*-weighted echo-planar images were obtained with a multiband sequence (axials co-planar with anterior-posterior commissure, 4 runs with 480 volumes and 60 slices per volume each, slice thickness = 2.5 mm, distance factor = 0%, FOV = 210 mm, voxel size = 2.5 mm isotropic, TR = 1000 ms, TE = 29 ms, flip angle = 56°, multiband acceleration factor = 4). Structural scans were acquired using a high-resolution T1-weighted magnetisation-prepared rapid-acquisition gradient echo (MPRAGE) sequence (sagittal orientation, slice thickness = 0.8 mm, distance factor = 50%, FOV = 260 mm, voxel size = 0.8 mm isotropic, TR = 1900 ms, TE = 2.54 ms, flip angle = 9°). We applied generalised autocalibrating partially parallel acquisitions (GRAPPA) with an acceleration factor of 2 for parallel imaging. T1 origin was subsequently reoriented to the anterior commissure to improve normalisation during preprocessing. Data were preprocessed and analysed using Statistical Parametric Mapping (SPM12; The Wellcome Centre for Human Neuroimaging, London, UK; https://www.fil.ion.ucl.ac.uk/spm/software/spm12) running on MATLAB R2017b (The MathWorks Inc., Natick MA, USA; https://www.mathworks.com/products/matlab.html). To allow for signal equilibration, the first four volumes of each functional scan were discarded as dummy volumes. Images were realigned to the mean functional image using a 6-parameter rigid body transformation, then coregistered to the individual structural scan, normalised to Montreal Neurological Institute (MNI) space with resampling to 3 mm isotropic voxel size, and finally spatially smoothed with a Gaussian kernel of 6 mm full-width at half-maximum. Successful coregistration and normalisation were verified through visual inspection.

### 2.5 Analysis approach

Behavioural data were processed and analysed in R (version 3.6.2; https://www.r-project.org). First, we checked for outliers and excluded runs with extreme values (above the third quartile plus three times the interquartile range (IQR) or below the first quartile minus three times the IQR; Kassambara et al., 2020). The few instances where we excluded data are described in the respective results sections. Next, we constructed linear mixed-effects models (LMEMs; Brauer & Curtin, 2018) based on the tutorial referenced in Singmann & Kellen (2019), using the afex package (Singmann et al., 2020). To confirm that stressor aversiveness and perceived control did not differ between conditions or over runs, we set up models with condition and run as fixed effects, including an interaction term. Given that condition and run were nested within participants, we incorporated by-participant random intercepts, random slopes for condition and run, and an interaction term to capture random variation. Models testing for condition-dependent differences in outcome measurements (stress and helplessness ratings, RTs, correct responses, BPM) comprised the same random effects. However, only condition was included as a fixed effect because we were not focused on time-dependent changes. For RTs and correct responses trial-by-trial data could be used. Hence, the latter was analysed using a generalised LMEM, appropriate for binary dependent variables (i.e., correct/incorrect). We began by fitting each model with the maximal random effects structure in order to minimise Type I error (Barr et al., 2013). We applied the “bobyqa” optimiser and set the number of model iterations to 10,000. If a model failed to converge or represented a singular fit, we pruned the random effects structure until the model converged without warnings. More precisely, we first removed correlations between random intercepts and random slopes, then random slopes themselves. All final models included participant-level random intercepts (see the supplement for details on final models). We verified that effects held across the maximal and reduced model. For post-hoc comparisons, we used the emmeans package (Lenth, 2020) and employed Holm-Bonferroni correction. Additionally, we applied Bonferroni correction to account for multiple dependent variables in our investigation of controllability-related effects. Only corrected significance levels are reported. We describe model details as suggested by Meteyard and Davies (2020) and state denominator degrees of freedom with decimals as recommended by Brauer & Curtin (2018).

With regard to the fMRI data, we followed SPM’s two-level general linear modelling approach to investigate effects of stress and controllability, focusing on the indicator (anticipation-phase) and subsequent fixation phase. We decided to disregard the feedback phase because the procedure employed to match stress durations across CON and UNCON trials resulted in slightly different trial structures making the feedback phase potentially less comparable. The first-level model comprised six regressors: indicator (CON, UNCON, baseline) and fixation (CON, UNCON, baseline), each modelled using the onset and length of the event (1500 ms and 4000 ms, respectively) convolved with the hemodynamic response function. Additionally, six parameters were included as nuisance regressors to account for any variance associated with motion (the Supplemental Figure visualises the model design matrix). The high pass filter cut-off period was set to 128 s Two contrasts were computed at the participant level, each for indicator and fixation: stress (CON + UNCON) versus baseline and CON versus UNCON. We also examined the inverse contrasts. The resulting beta images were then subjected to a second-level flexible factorial model comprising condition and the individual participant factor. Main effects of stress were assessed at the whole-brain level. Family-wise error (FWE) correction was performed for voxel-level inference at a threshold of alpha = 0.01 and a cluster-extent threshold of 10 voxels. Controllability effects were investigated in the same manner, however, for the fixation phase, we restricted analyses to our a priori defined regions of interest (ROIs) – the vmPFC and the bilateral amygdala (SVC = small volume correction). The vmPFC mask was taken from Bhanji et al. (Supplement 1 from Delgado et al., 2016) and the left and right amygdala mask was created based on the Harvard-Oxford brain atlas (Desikan et al., 2006), including only voxels with at least a 25% tissue probability. For the ROI analysis, FWE-correction was performed for voxel-level inference using a threshold alpha = 0.05. Peak voxel locations were labelled using the automated anatomical labelling toolbox (AAL3; Rolls et al., 2020).

In light of the animal research findings on controllability-dependent connections between the vmPFC, DRN and its output-regions (e.g., amygdala), we also examined differences in vmPFC connectivity under CON versus UNCON (see Supplement 13 for details).

Finally, we extracted parameter estimates from the vmPFC region more activated under CON compared to UNCON. Specifically, we output beta weights for baseline, CON, and UNCON regressors for each participant and each run. Firstly, this served to more thoroughly address what might be driving the observed effect, i.e., differences in activation or deactivation or possibly a mixture of both. Secondly, it enabled us to examine links between vmPFC activation and ratings (stress and helplessness) for each condition separately. To this end, mean centered beta weights and an interaction term of beta weight and condition were added as fixed effects to the previously constructed LMEMs investigating stress and helplessness ratings.

Behavioural data and analysis code is available on the Open Science Framework: https://osf.io/8qpme/.

## 3. Results

### 3.1 Manipulation check

First, we compared average stress durations between CON and UNCON conditions. A *t*-test indicated imperfect yoking: participants had experienced on average slightly longer stress phases in CON compared to UNCON trials (*t*_44_ = 6.33, *p* < .0001, *d* = 1, CON-UNCON stress duration difference: *M* = 1.73 s, *SD* = 1.83). Two participants reported receiving more electric stimuli in CON compared to UNCON trials, effectively noticing this imbalance. They were thus excluded from analysis. To account for the mismatch and its varying magnitude between participants, we included random slopes for the CON-UNCON stress duration difference in subsequent LMEMs. All results remained unchanged. We assessed stressor aversiveness ratings (Figure 2a) and observed no differences between conditions, participants generally rated the electric stimuli as aversive (global *M* = 63.75; *F*_1, 264.15_ = 2.16, *p* = .143). There was a small decrease in ratings over runs, indicating slight habituation to the stressor (*F*_1, 44.12_ = 11.66, *p* = .001). No interaction of condition x run was indicated (*F*_1, 264.15_ = 0.53, *p* = .469). In line with our manipulation, participants perceived significantly more control in CON compared to UNCON trials (*F*_1, 44.24_ = 104.04, *p* < .0001). These differences became more pronounced over runs with ratings increasing for CON, but decreasing for UNCON (*F*_1, 217.75_ = 6.08, *p* = .014; Figure 2b). Detailed model results can be found in the supplementary material (Supplemental Tables 1 and 2).

**Figure 2.**
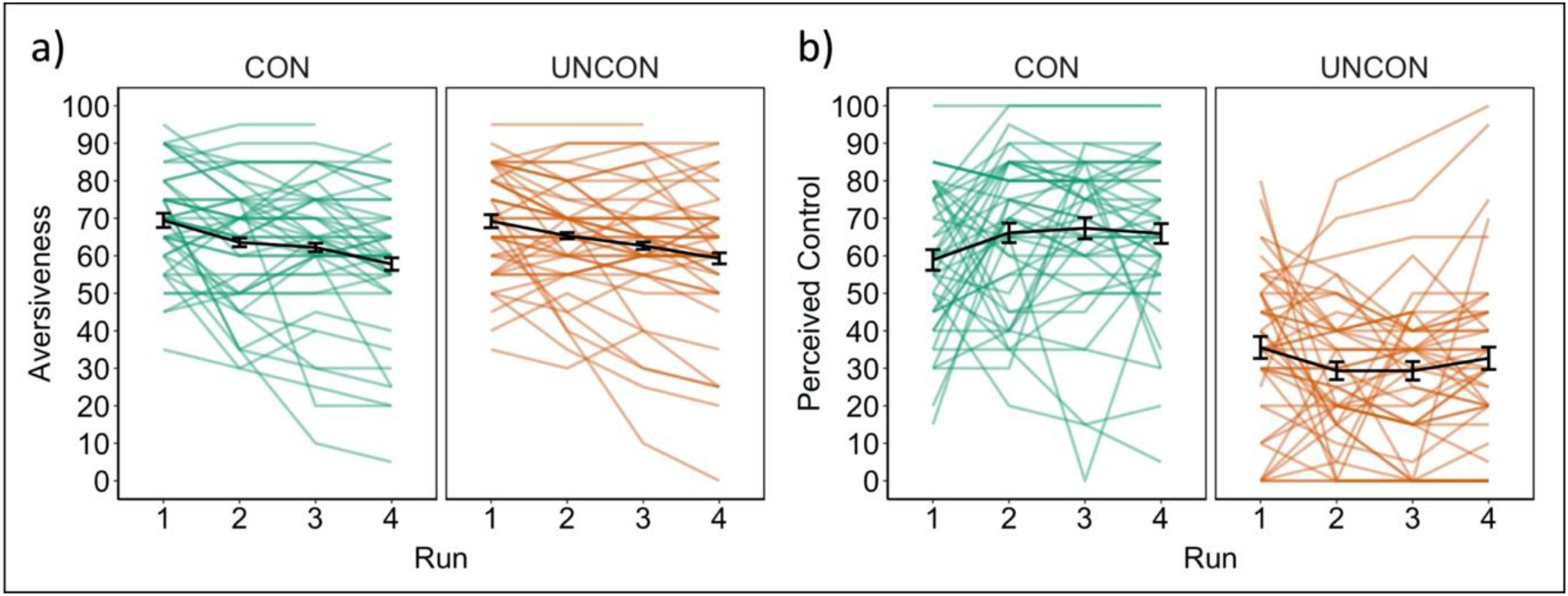
Manipulation checks: (a) across conditions, participants rated the stressor as very aversive with only slight habituation over runs. (b) They reported perceiving significantly more control for controllable stress (CON) trials compared to uncontrollable stress (UNCON) trials. This effect became slightly more pronounced over runs. Error bars denote within-subject standard errors.

### 3.2 Ratings, RTs, accuracy, and heart rate

Participants’ stress ratings were significantly higher for stress compared to baseline trials (*F*_2,44.12_ = 36.41, *p* < .0001; CON vs. baseline and UNCON vs. baseline: *p* < .0001), however, there was no difference between CON and UNCON (*p* = .423; Figure 3a). As expected, helplessness ratings were lower for CON compared to UNCON (*F*_1, 40.65_ = 34.03, *p* = .001; Figure 3b). For the analysis of RTs, one run from one participant was excluded as an outlier due to extremely short RTs. Participants responded faster in CON compared to both UNCON and baseline trials (*F*_2, 4743.88_ = 84.02, *p* < .0001; CON vs. UNCON: *p* = .001 and CON vs. baseline: *p* < .0001). RTs were slightly shorter under UNCON compared to baseline (*p* = .014; Figure 3c). Concerning correct responses, data from one participant and one run each from four other participants were excluded from analysis because they demonstrated extremely low performance. In general, participants showed high accuracy (*M* = 0.87, *SD* = 0.14), but differences across conditions emerged (*X*^2^ = 27.96, *df* = 2, *p* < .0001; Figure 3d). Performance was significantly better under stress compared to baseline (CON vs. baseline: *p* < .0001; UNCON vs. baseline: *p* = .007). Accuracy was slightly higher in CON compared to UNCON trials (*p* = .037). In terms of heart rate, data from one participant and one run each from two other participants were excluded from analysis because their BPM represented extreme deviations from the sample mean, probably reflecting recording equipment issues. Participant’s heart rate differed across conditions (*F*_2, 375.83_ = 7.01, *p* = .001; Figure 3e). It was significantly higher under CON compared to baseline (*p* = .001), though differences in absolute values were small. No differences in BPM were observed between CON and UNCON or UNCON and baseline (both contrasts: *p* = .580). The supplementary material contains detailed model results (Supplemental Tables 3-7).

**Figure 3.**
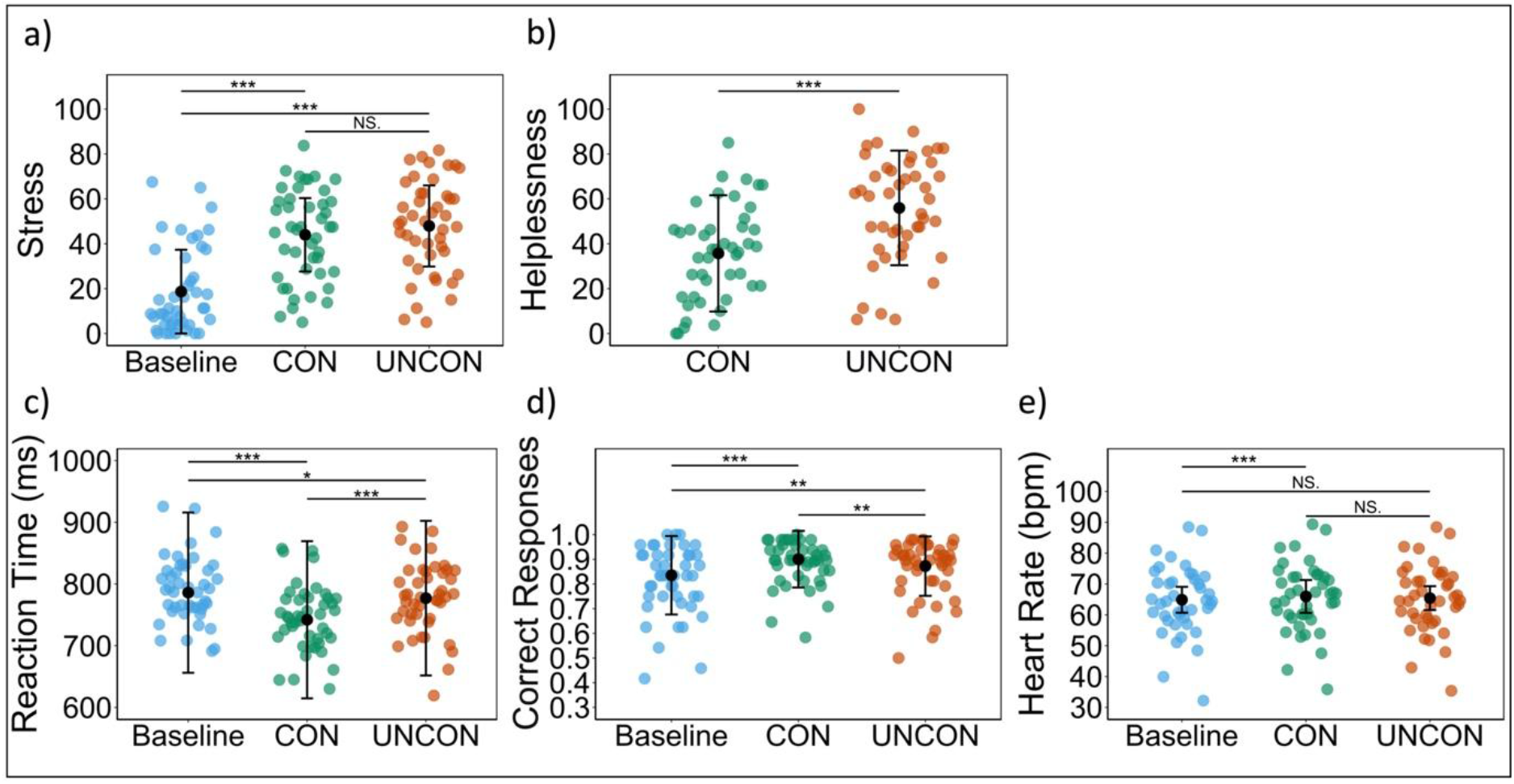
Ratings, RTs, accuracy, and heart rate: (a) subjectively rated stress levels were higher for stress trials compared to baseline. (b) Participants reported feeling less helpless under controllable stress (CON) compared to uncontrollable stress (UNCON). (c) Reaction times were fastest in controllable trials, followed by uncontrollable trials, then baseline. (d) accuracy was generally highest in controllable trials, followed by uncontrollable trials, then baseline. (e) Heart rates were higher under controllable stress compared to baseline, but no differences emerged between controllable and uncontrollable trials. Error bars denote within-subject standard errors.

### 3.3 Neuroimaging results

To examine effects of stress during anticipation and fixation, we contrasted stress (CON+UNCON) and baseline trials. During fixation we observed increased activation in e.g., the anterior and posterior insula (peak voxel: −32 24 8, Z > 8, *p*_FWE_ROI_ < .001, 2313 voxels and −34 −20 16, Z > 8, *p*_FWE_ROI_ < .001, 362 voxels, respectively; Figure 4a), the left postcentral gyrus, and the supplementary motor area (SMA; Figure 4b; see Table 1 for all clusters). Areas less activated under stress compared to baseline included the right paracentral lobule (Figure 4c), the right postcentral gyrus, the right inferior occipital gyrus, the left fusiform gyrus (Figure 4d), and the hippocampus among others (Table 1). During anticipation, similar clusters of significant BOLD response changes emerged (Supplemental Table 9).

**Figure 4.**
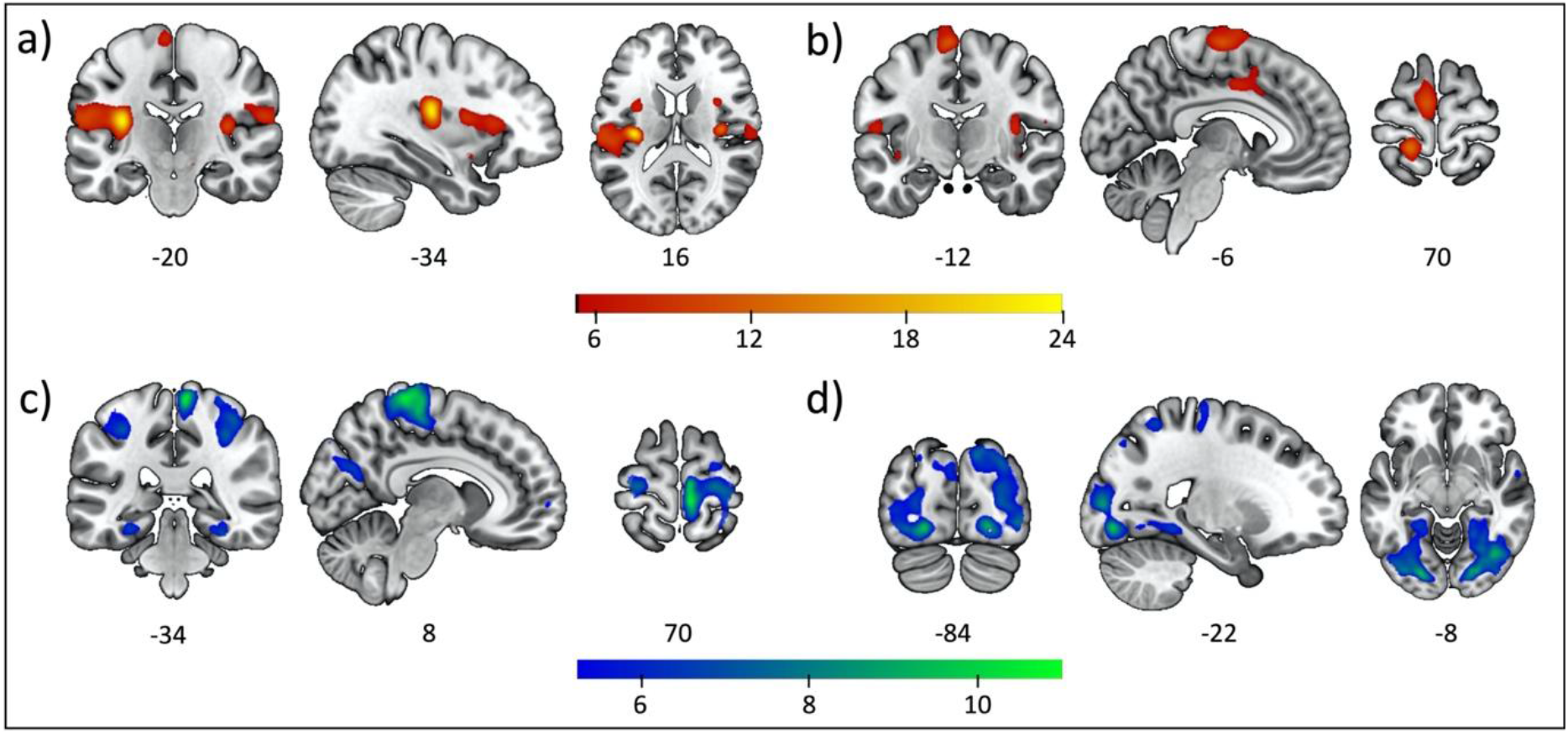
Stress effects: the differential contrast Stress > Baseline revealed activation in e.g., (a) the bilateral insula, (b) supplementary motor area and premotor cortex. The inverse contrast (Baseline > Stress) yielded significant clusters in e.g., (c) the paracentral lobule, (d) fusiform and parahippocampal gyrus. Thresholded T-maps overlaid with the MNI 152 template.

**Table 1.**
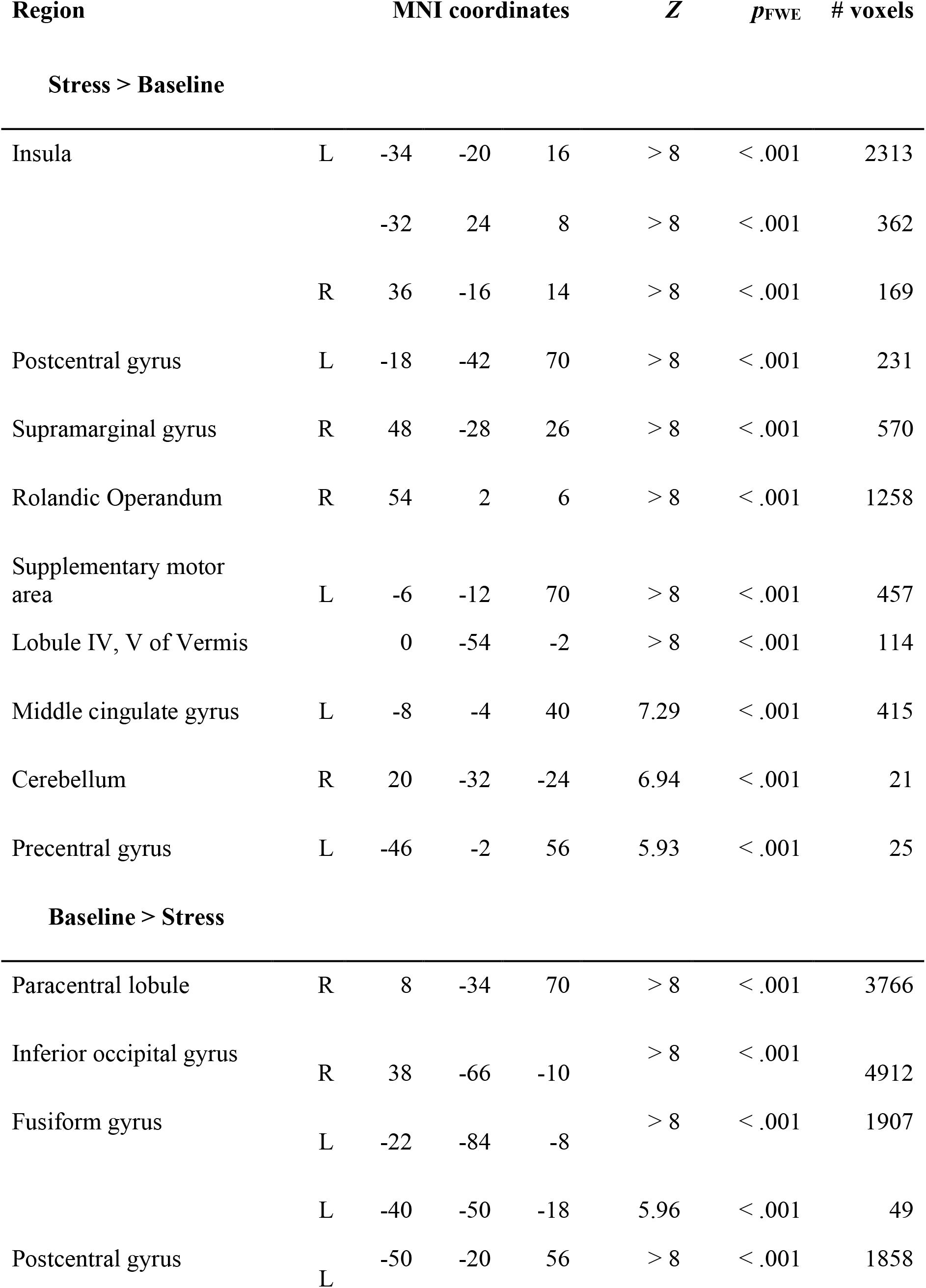
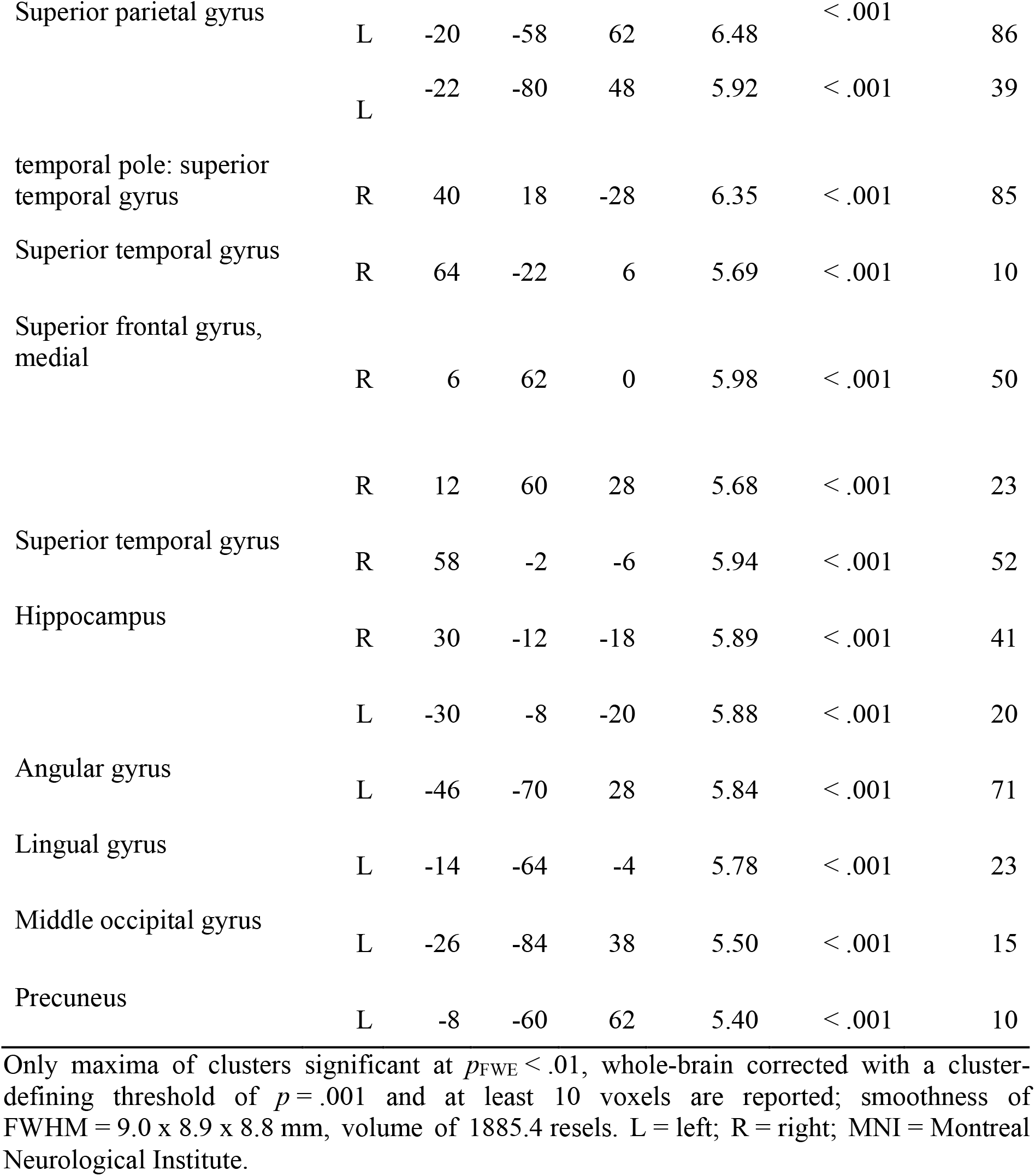
Brain activations associated with stress during fixation

To investigate effects of controllability during fixation, we contrasted CON trials with UNCON trials and restricted our analyses to our ROIs, the vmPFC and the amygdala. The CON > UNCON contrast yielded a significant cluster in the left vmPFC with two local maxima (peak voxel: −6 36 −8, Z = 4.78, *p*_FWE_ROI_ = .004 and −12 44 −6, Z = 4.68, *p*_FWE_ROI_ = .007, respectively, 169 voxels; Figure 5a; Figure 6a). No significant clusters emerged in the bilateral amygdala. An additional exploratory whole-brain analysis revealed no sizeable suprathreshold clusters. The inverse contrast indicated less activation of stress-activated areas, i.e., the insula (Figure 5b), as well as the cingulate gyrus, hippocampus, fusiform gyrus, motor and visual cortices under CON compared to UNCON (Table 2). A ROI analysis restricted to the bilateral amygdala also revealed a significant cluster with two local maxima in the right amygdala (peak voxel: 26 −2 −18, Z = 3.69, *p*_FWE_ROI_ = .014 and 28 2 −20, Z = 3.56, *p*_FWE_ROI_ = .021, respectively, 15 voxels) and a smaller cluster in the left amygdala (peak voxel: −24 −4 −14, Z = 3.25, *p*_FWE_ROI_ = .048, 1 voxel). These results remained unchanged after we included the stress duration as a covariate at the second level.

**Figure 5.**
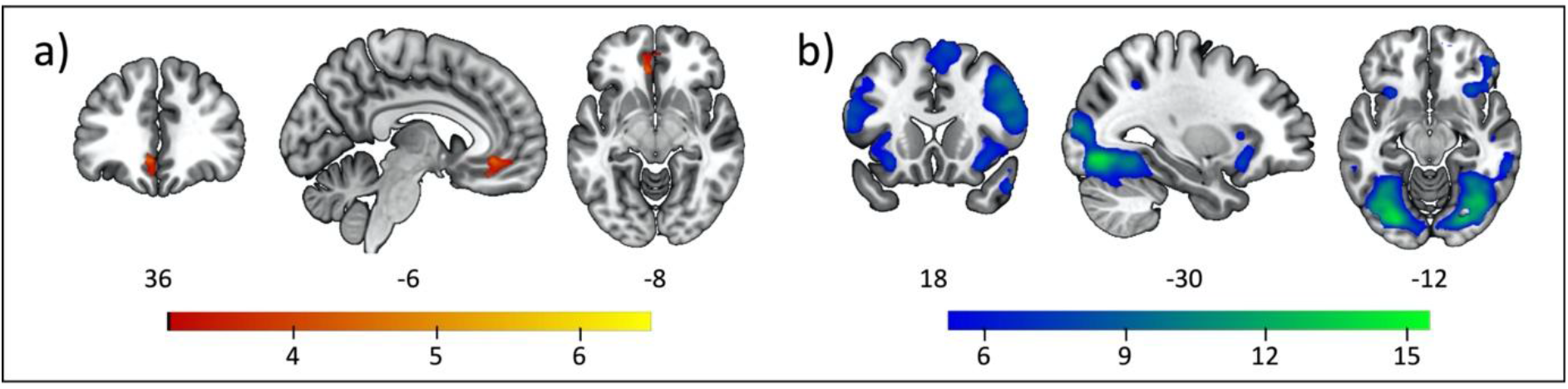
Stressor controllability effects: region of interest analysis contrasting controllable stress (CON) and uncontrollable stress (UNCON) trials revealed activation in (a) the vmPFC. The inverse contrast (UNCON > CON) demonstrated decreased activation in stress-responsive areas, e.g., (b) insula. Thresholded T-maps overlaid with the MNI 152 template.

**Figure 6.**
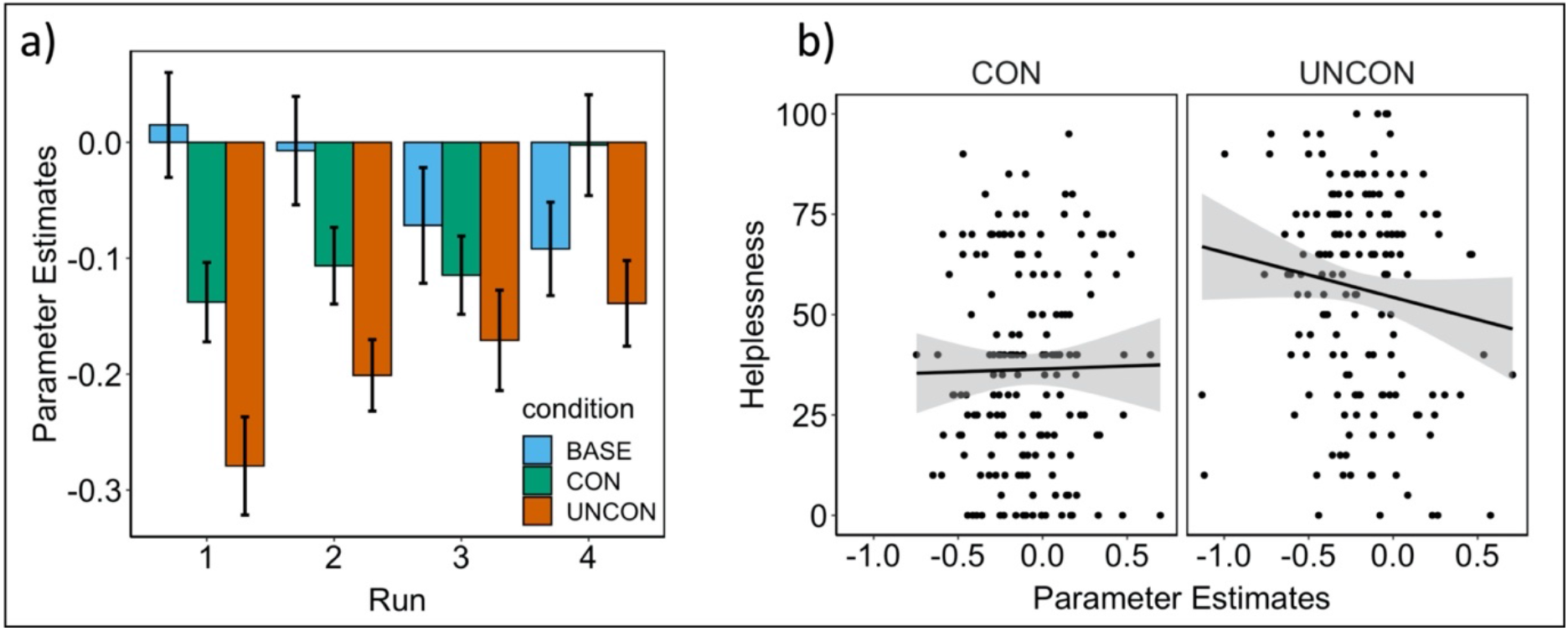
(a) Parameter estimates for baseline (BASE), controllable stress (CON), and uncontrollable stress (UNCON) extracted from the vmPFC cluster. Error bars denote within-subject standard errors. (b) VmPFC activation modulates helplessness ratings for UNCON, but not CON. One data point per participant and run in each condition (as used for mixed-effects modelling).

**Table 2.**
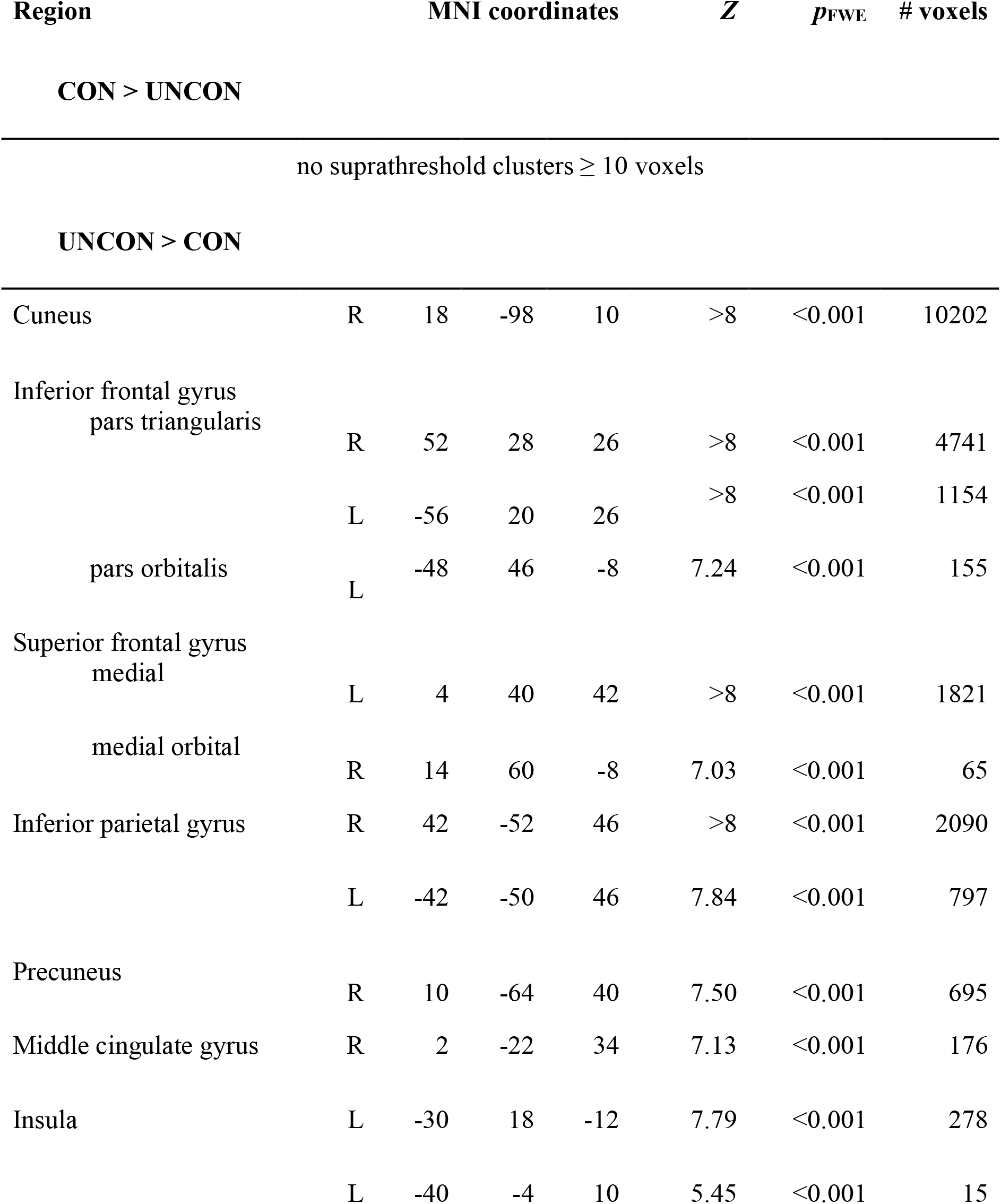
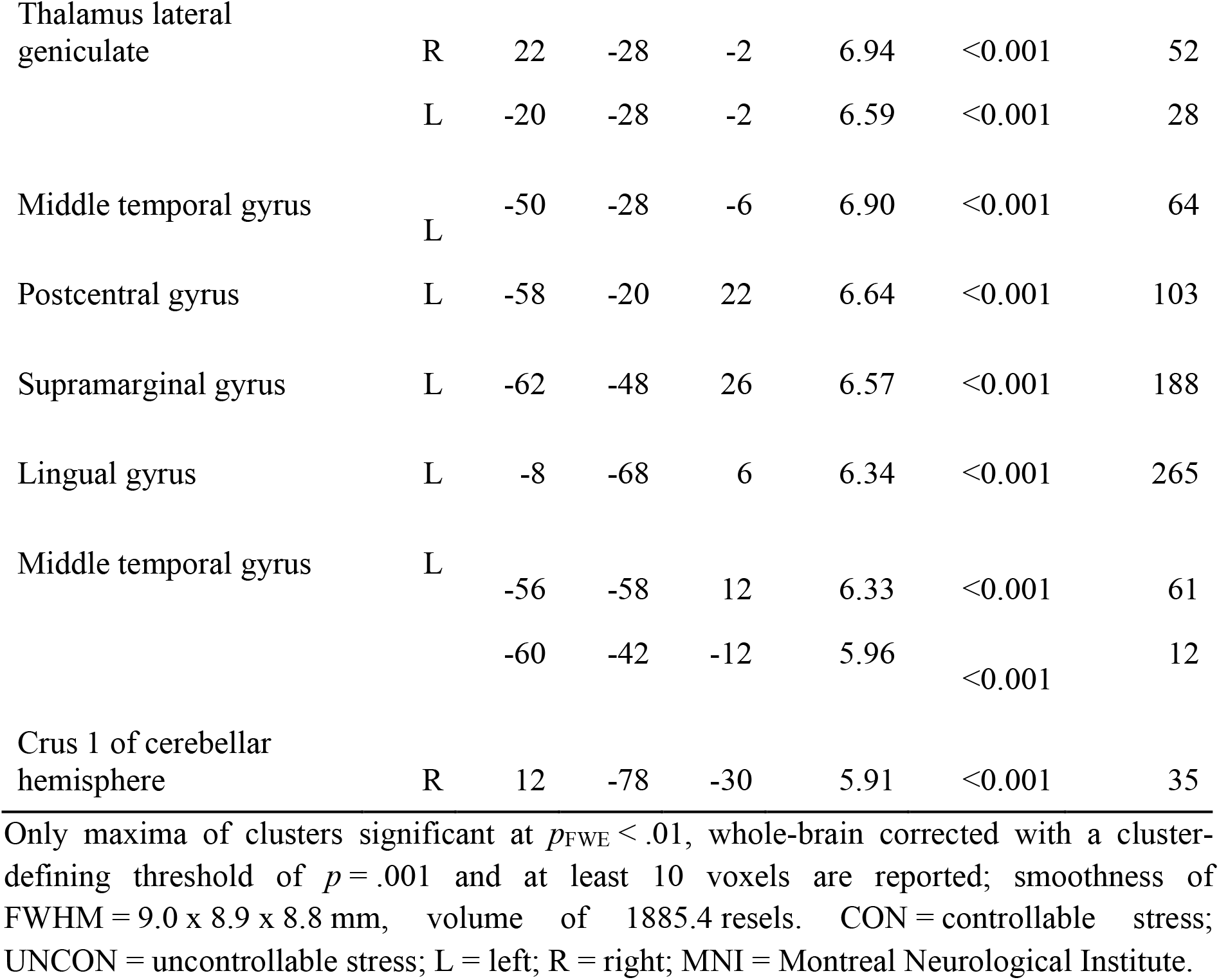
Brain activations related to stressor controllability during fixation

| Region |  | MNI coordinates |  |  | *Z* | *p*<sub>FWE</sub> | # voxels |
| --- | --- | --- | --- | --- | --- | --- | --- |
| CON > UNCON |  |  |  |  |  |  |  |
| no suprathreshold clusters ≥ 10 voxels |  |  |  |  |  |  |  |
| UNCON > CON |  |  |  |  |  |  |  |
| Cuneus | R | 18 | -98 | 10 | >8 | <0.001 | 10202 |
| Inferior frontal gyrus<br>pars triangularis | R | 52 | 28 | 26 | >8 | <0.001 | 4741 |
|  | L | -56 | 20 | 26 | >8 | <0.001 | 1154 |
| pars orbitalis | L | -48 | 46 | -8 | 7.24 | <0.001 | 155 |
| Superior frontal gyrus<br>medial | L | 4 | 40 | 42 | >8 | <0.001 | 1821 |
| medial orbital | R | 14 | 60 | -8 | 7.03 | <0.001 | 65 |
| Inferior parietal gyrus | R | 42 | -52 | 46 | >8 | <0.001 | 2090 |
|  | L | -42 | -50 | 46 | 7.84 | <0.001 | 797 |
| Precuneus | R | 10 | -64 | 40 | 7.50 | <0.001 | 695 |
| Middle cingulate gyrus | R | 2 | -22 | 34 | 7.13 | <0.001 | 176 |
| Insula | L | -30 | 18 | -12 | 7.79 | <0.001 | 278 |
|  | L | -40 | -4 | 10 | 5.45 | <0.001 | 15 |
| Thalamus lateral geniculate | R | 22 | -28 | -2 | 6.94 | <0.001 | 52 |
|  | L | -20 | -28 | -2 | 6.59 | <0.001 | 28 |
| Middle temporal gyrus | L | -50 | -28 | -6 | 6.90 | <0.001 | 64 |
| Postcentral gyrus | L | -58 | -20 | 22 | 6.64 | <0.001 | 103 |
| Supramarginal gyrus | L | -62 | -48 | 26 | 6.57 | <0.001 | 188 |
| Lingual gyrus | L | -8 | -68 | 6 | 6.34 | <0.001 | 265 |
| Middle temporal gyrus | L | -56 | -58 | 12 | 6.33 | <0.001 | 61 |
|  |  | -60 | -42 | -12 | 5.96 | <0.001 | 12 |
| Crus 1 of cerebellar hemisphere | R | 12 | -78 | -30 | 5.91 | <0.001 | 35 |

Only maxima of clusters significant at $p_{\text{FWE}} < .01$ , whole-brain corrected with a cluster-defining threshold of $p = .001$ and at least 10 voxels are reported; smoothness of FWHM = 9.0 x 8.9 x 8.8 mm, volume of 1885.4 resels. CON = controllable stress; UNCON = uncontrollable stress; L = left; R = right; MNI = Montreal Neurological Institute.

During anticipation, for CON > UNCON, significant clusters emerged in the inferior frontal, middle temporal, and middle frontal gyrus (Supplemental Table 10). Regions that were less activated under anticipation of control compared to lack of control included the inferior occipital and lingual gyrus as well as a vmPFC area overlapping with the cluster that had emerged from the CON > UNCON contrast during fixation. The analysis of vmPFC connectivity yielded no suprathreshold clusters, neither at the whole-brain level nor when restricting the analysis to the cluster in the right or left amygdala that had emerged in the GLM analysis for the UNCON>CON contrast.

Parameter estimates extracted from the vmPFC cluster reflected a general deactivation of this region under acute stress, which was further supported by an additional ROI analysis comparing neural activation at baseline vs stress during fixation (Supplemental Table 11). Interestingly, the vmPFC parameter estimates were differentially associated with helplessness ratings for UNCON and CON, as evidenced by a significant condition x beta weight interaction (*F*_1, 296.20_ = 7.89, *p* = .005; CON vs. UNCON slopes: *p* = .005; Figure 6b; model details in Supplemental Table 8). Specifically, higher beta weights were associated with lower helplessness ratings for UNCON, whereas no such modulation was observed under CON. No interaction effect emerged for stress ratings (*F*_2, 387.48_ = 1.97, *p* = .14).

## 4. Discussion

The present study used a translational experimental design drawing upon animal research on learned helplessness in order to investigate immediate differential effects of stressor controllability (controllable vs. uncontrollable stress) on neural activation patterns, subjective ratings of stress and feelings of helplessness, RTs and task performance as well as heart rate. In line with our hypotheses, we observed more vmPFC recruitment and lower helplessness ratings under controllable compared to uncontrollable stress. Both conditions were rated similarly stressful compared to baseline, however, contrary to our hypothesis, no differences between controllable and uncontrollable stress emerged. We also noted differences in RTs and performance that were not anticipated. Specifically, participants responded faster and more accurately under stress compared to baseline. The finding that RTs were shortest and accuracy highest under controllable compared to uncontrollable stress was, however, in line with our hypotheses. In terms of heart rate, we had expected more BPM under stress compared to baseline, and an even more elevated heart rate under uncontrollable than controllable stress. Contrary to our hypothesis, we recorded more BPM under controllable stress compared to baseline and observed no other significant contrasts. In the following, we will first discuss the behavioural and heart rate results and then the neuroimaging results in more detail.

### 4.1 Behavioural and heart rate results

As anticipated, ratings indicated greater perceived control for controllable compared to uncontrollable stress, which confirms that our experimental manipulation aligned with participants’ subjective experience. This might seem trivial as the objective controllability was explicitly signalled at the beginning of each trial. However, we still observed inter-individual variation in subjectively rated controllability, suggesting that – as in previous between-group designs (Meine et al., 2020; Wanke & Schwabe, 2020a) – objective and subjectively experienced controllability does not match in each individual for every trial. Furthermore, ratings of perceived control for controllable trials were generally not at ceiling, likely reflecting the fact that participants had to endure a certain amount of stress even in these trials. This was a necessary constraint to allow the investigation of direct effects of stressor controllability under acute stress. Feelings of helplessness were correspondingly higher for uncontrollable compared to controllable stress, replicating effects from our earlier studies (Meine et al., 2020). Since learned helplessness is commonly discussed as a model for depression (Pryce et al., 2011; Maier & Seligmann, 2016), the evident modulation of helplessness by controllability bears significant potential for clinical intervention.

Participants reported feeling more stressed in trials with aversive stimulation, but they showed concurrently faster RTs and greater task accuracy compared to baseline. Whereas the ratings are partly in line with our hypotheses and reflect successful stress induction, we had expected stress-related RT and performance decrements. However, given the stressfulness of the aversive stimulation, the results can easily be explained by the motivation to terminate the stressor. Baseline trials offered no such pressing need for correct responses. Previous research has shown some beneficial effects of stress on concentration (Degroote et al., 2020) and performance (Beste et al., 2013; Travis et al., 2020), particularly under low task demands (Kim et al., 2017). Indeed, our task was quite simple, as confirmed by the generally high rate of correct responses. Control over the stressor evidently served as further encouragement to perform well. As expected, participants responded more quickly and accurately when their response was instrumental in terminating the stressor than when aversive stimulation was uncontrollable. Surprisingly, performance was better under uncontrollable stress than in the baseline condition. This was unexpected because participants knew that their responses had no consequence in uncontrollable trials. However, since accurate and fast responses were equally inconsequential in baseline trials (where no stressor was applied), two explanations might be considered for these findings. First, participants may just have complied with the instruction to respond as quickly and accurately as possible in all conditions and, second, they may have harboured hope that their responses might show some effect after all, especially when subjected to uncontrollable stress.

Contrary to our hypothesis and despite the apparent effectiveness of our manipulation, no robust differences in heart rate emerged. Recordings showed only slightly higher heart rate under controllable stress compared to baseline. Yet, a large body of research has described elevated heart rates under acute stress (Hellhammer & Schubert, 2012; Orem et al., 2019; Sandner et al., 2020). Since we employed an event-related design with relatively short trials, this setup may have prevented a full return to baseline heart rate levels between conditions. Hence, contrasts may have been confounded by carry-over effects and should be interpreted with caution. Possibly, the analysis of heart rate variability – an increasingly popular stress marker (Thayer et al, 2012) – could offer more fine-grained results, but this generally requires electrocardiogram recordings (Schäfer & Vagedes, 2013). Furthermore, guidelines advocate durations of at least five minutes for short-term assessments (Malik et al., 1996). Unfortunately, this suggests that our data do not yield a reliable assessment of differences in heart rate, however, this was not our main focus. Instead, our main objective was to unravel neurobiological differences underlying stressor controllability.

### 4.2 Neuroimaging results

In line with our hypotheses, results demonstrate stressor-induced activation in brain regions commonly implicated in the stress response, e.g., the insula (Van Oort et al., 2017; Sandner et al., 2020). Research indicates functionally diverse roles of the anterior and posterior insula, ascribing roles in affective- and pain processing, respectively (Singer et al., 2004). For instance, paradigms employing negative feedback to induce stress have reported corresponding activations in the anterior insula in particular (Sandner et al., 2020). Following the principles of translational research, we employed a physical stressor akin to animal paradigms. However, we found both posterior and anterior parts of the insular cortex to be activated in response to this stressor. In general, human stress research spans many different types of stressors, e.g., electric stimuli (Hartley et al., 2014), thermal stimulation (Bräscher et al., 2016), loud sounds (Henderson et al., 2012), video clips of snakes (Kerr et al., 2012), and unsolvable reasoning tasks (Bauer et al., 2003). Furthermore, research has shown stress-related deactivations of frontal areas (Arnsten, 2015) and the limbic system (Pruessner et al., 2008). Our finding that the hippocampus was less activated under stress compared to baseline is consistent with this literature.

Critically, we observed the expected controllability-dependent differences in vmPFC activation. More precisely, vmPFC recruitment under stress was greater when participants knew they could end the aversive stimulation compared to when there was no contingency between their response and stressor termination. As stress was shown to suppress cortical regions involved in higher-order cognition, i.e, the vmPFC (Arnsten, 2015), the analysis of parameter estimates extracted from the vmPFC helped clarify what was driving the significant contrast. The vmPFC was in fact deactivated both during controllable and uncontrollable stress, but considerably more so in the latter condition. Hence, we observed a controllability-dependent difference in the magnitude of deactivation rather than activation. This finding lends further support to the notion that uncontrollable stress in particular presents a powerful switch which effectively shuts down the PFC (Datta & Arnsten, 2019). Instrumental control, on the other hand, is deemed a protective factor that attenuates stress effects (Henderson et al., 2012; Hartley et al., 2014; Maier & Seligman, 2016). In line with this, control was associated with decreased activation of the amygdala as well as deactivation of the insula, postcentral and middle cingulate gyrus – regions responsive to stress as identified in our analysis. Many of these areas have previously been found to be modulated by stressor controllability. Indeed, Wang and Delgado (2021) observed lower amygdala activation under exposure to controllable aversive tones compared to uncontrollable tones and reported cluster coordinates comparable to ours. Similarly, another very recent study noted decreased activation in the insula and other threat-related areas under controllable aversive electric shocks (Limbachia et al., 2021).

Contrary to our hypothesis, we observed no differences in vmPFC connectivity between controllable and uncontrollable stress. The overall down-regulatory effect that stress exerts on the vmPFC may well have prevented putative differences from reaching suprathreshold levels. In fact, studies that have found the expected condition-dependent connectivity patterns in humans have largely focused on anticipatory responses rather than investigating blood-oxygen-level-dependent (BOLD) signalling under acute stress (Kerr et al., 2012; Wanke & Schwabe, 2020b). Because the fixation phase in our design captures participants’ neural responses just before they were prompted to terminate the stressor, it effectively also constitutes an anticipatory phase, albeit under acute stress. Interestingly, during the presentation of the indicator (i.e., anticipation without stress), participants showed more activation in the vmPFC when an uncontrollable trial was signalled, but under stress they subsequently displayed enhanced vmPFC recruitment when they knew they could terminate the stressor. Although the former finding replicates results by Wanke & Schwabe (2020b), the apparent reversal in vmPFC recruitment induced by aversive stimulation encourages further research into timing-dependent effects.

Taken together, our results are consistent with animal research in which the vmPFC was shown to downregulate the stress response upon detecting control (Maier & Seligman, 2016). If the neural mechanisms by which controllability modulates our reaction to and appraisal of stressful events are parallel to those found in animals, translational research may offer great promise for the treatment of stress-related mental dysfunction. Animal research has already begun to test pharmacological interventions aimed at enhancing vmPFC activation to ameliorate stress effects. Although these studies can boast considerable successes (Amat et al., 2016), it seems important to consider and pursue further non-pharmacological treatment routes. Alongside behavioural interventions, advances in neuroimaging are starting to open up opportunities for non-pharmacological clinical interventions, such as neurofeedback. In a recent study, Keynan et al. (2019) employed simultaneous electroencephalography (EEG)-fMRI to individually identify signals from the amygdala in military personnel. Using only EEG, they subsequently targeted this so-called amygdala electrical fingerprint in neurofeedback sessions and trained participants to deliberately downregulate activation. Although further validation may be required, a similar approach can be envisaged in clinical contexts. A recent finding showing that vmPFC activation varies with subjective stress ratings (Orem et al., 2019) supports this idea. In this study, we did not observe a relationship between vmPFC recruitment and stress ratings. However, we could show that increased vmPFC recruitment was associated with reduced feelings of helplessness under lack of control. Since helplessness has been heavily implicated in the aetiology of depression (Pryce et al. 2011), this finding represents a critical contribution to the literature. Perhaps, modulations of vmPFC activation are particularly beneficial in uncontrollable contexts. VmPFC recruitment was already shown to be predictive of recovered active avoidance behaviour following passivity induced by uncontrollable stress (Wang & Delgado, 2021). Future research should further investigate the link between vmPFC activation and coping behaviours.

In general, interventions must be directed not only at modulating stress responses on the neurophysiological level, but also in terms of subjective experience. After all, the self-report necessarily describes the symptoms most relevant to the patient and can readily be assessed by clinicians. Moreover, research has highlighted the importance of subjectively perceived controllability over objectively given control. Wanke and Schwabe (2020a) found impaired working memory processes to be associated specifically with perceived lack of control over an aversive stimulus rather than objective control or even stress per se. Similarly, Hancock & Bryant (2018a; 2018b) reported links between controllability expectations and stress-avoidance, especially in patients with posttraumatic stress disorder (PTSD).

### 4.3 Limitations and future directions

The present study has some limitations that must be considered when interpreting the results.

First, the within-subject nature of the design led us to explicitly indicate the different trial types and instruct participants as to the correct responses. This helped participants capitalise on the opportunity to terminate the stressor in controllable trials, in line with our intended manipulation. Hence, we were able to clearly differentiate controllable and uncontrollable stress trials from the start of the experiment, but at a cost to our translational efforts. In the original animal experiments, the subjects are required to figure out by themselves how to terminate aversive stimulation. Furthermore, as discussed in the introduction, the experience of control over a stressor may affect responses to subsequent uncontrollable events – and vice versa (Maier & Seligman, 2016). Moreover, any within-subject design is potentially vulnerable to carry-over effects. To prevent such problems, we set up a trial sequence optimised for detecting condition-dependent differences in BOLD signal. The explicitly indicated constant change in trial type also precluded the establishment of an enduring sense of control or feeling of helplessness. In fact, the ratings of perceived control for controllable and uncontrollable trials became more differentiated over time (Figure 2b) and parameter estimates extracted from the vmPFC likewise showed no evidence of overlapping controllability effects (Figure 6a). After all, we were not interested in controllability-dependent effects on measurements following stress exposure, but rather examined the immediate effects of stressor controllability. For this reason, we also refrained from measuring cortisol – an indicator of the stress response characterised by a rather slow activation profile. Ratings, RTs, and task performance should not have been significantly influenced by the event-related experiment structure, but differences in heart rate may have been clearer in a block design. However, our main focus was on detecting differences in neural activation depending on stressor controllability. Expecting stronger contrasts, we deliberately chose the within-subject design because the same participant could experience all conditions. Furthermore, we anticipated that the variation across trials would increase motivation and compliance. In addition, the smaller sample size required allowed more rapid testing and restricted the number of participants that had to undergo this rather uncomfortable experience.

Second, our experimental manipulation of control resulted in imperfect yoking of stress durations. Specifically, participants suffered slightly longer aversive stimulation in the controllable compared to the uncontrollable condition. Nevertheless, most participants noticed no difference or even reported receiving more electric stimuli in the uncontrollable condition. To account for the mismatch, we included the CON-UNCON stress duration difference in our LMEMs and re-ran relevant fMRI analyses, adding it as a covariate at the second level. If our manipulation is to be used in future studies, this slight imbalance demands improvements to our yoking algorithm. Still, it does not diminish the validity of our results. Rather, the fact that the vmPFC was less deactivated in controllable trials, even though stress phases were, on average, slightly longer than in uncontrollable trials, highlights the robustness of the reported condition-dependent neural activation patterns.

Third, the lack of a jittered time interval between indicator and fixation in our design precludes a clean differentiation of these two phases in terms of BOLD signal. However, results point to marked differences between phases. In response to the indicator announcing uncontrollable rather than controllable stress, the vmPFC was more activated. In contrast, under acute stress, vmPFC recruitment was greater for controllable compared to uncontrollable trials.

Fourth, this study investigated a rather homogeneous sample of healthy, well-educated young participants and results may not be generalised to the whole population. Follow-up investigations focusing on inter-individual differences could further validate controllability-dependent neural processing as a critical mechanism in human stress processing. For instance, the assessment of participants stratified for resilient outcome (Chmitorz et al., 2020) could provide important insights to fostering mental health.

## 5. Conclusion

This study provides further evidence that controllability in anticipation of as well as under acute stress is associated with distinct neural processes. Results largely corroborate animal findings in a human sample: we showed that the vmPFC was particularly involved in instrumental control and that its activation under stress was associated with a more favourable response. Specifically, our results indicate that vmPFC recruitment under uncontrollable stress attenuated feelings of helplessness. These findings encourage further research into targeted interventions that can promote mental health and resilience in the face of stress.

## Supporting Information

Please see the supplementary materials.

## Competing Interests

The authors declare no conflict of interest.

## Author Contributions

Conceptualisation, M.W.; methodology, L.E.M., B.M. and M.W.; software, L.E.M.; formal analysis, L.E.M., J.M. and B.M.; investigation, L.E.M. and J.M.; data curation, L.E.M. and J.M.; writing-original draft preparation, L.E.M.; writing-review and editing, L.E.M., J.M., B.M. and M.W.; visualisation, L.E.M. and J.M.; supervision, M.W.; project administration, L.E.M. and M.W.; funding acquisition, M.W.

## Data Accessibility

Behavioural data and analysis code will be made available on the Open Science Framework: https://osf.io/8qpme/ and will be live upon publication. Due to data protection issues, the MRI data cannot be made available to the public, but we have uploaded our code. The experiment code is available upon reasonable request from the corresponding author.

## Acknowledgements

This research was funded by the German Research Foundation (DFG), Collaborative Research Center 1193, Project C07. The funders had no role in the design of the study; in the collection, analyses, or interpretation of data; in the writing of the manuscript, or in the decision to publish the results. We thank all members of Project C07 for their vital contribution, and all participants for taking part in the study. Special thanks go to Maximilian Zerbe for help with coding the experimental paradigm and Heike Schmidt-Harth for help with data acquisition.

## Abbreviations

AAL: Automated anatomical labelling
BOLD: Blood-oxygen-level-dependent
BPM: Beats per minute
CON: Controllable stress
DRN: Dorsal raphe nucleus
DSM: Diagnostic and Statistical Manual of Mental Disorders
EEG: Electroencephalography
fMRI: Functional magnetic resonance imaging
FOV: Field of view
FWE: Family-wise error
GRAPPA: generalised autocalibrating partially parallel acquisitions
ITI: Inter-trial interval
IQR: Interquartile range
LMEM: Linear mixed-effects model
MPRAGE: magnetisation-prepared rapid-acquisition gradient echo
PTSD: posttraumatic stress disorder
ROI: Region of interest
RT: Reaction times
SCID: Structured clinical interview for DSM-IV
SMA: Supplementary motor area
SPM: Statistical parametric mapping
SVC: Small volume correction
TE: Echo time
TR: Repetition time
UNCON: Uncontrollable stress
vmPFC: Ventromedial prefrontal cortex

## Notes

### Competing Interest Statement

The authors have declared no competing interest.

