## Supplement for "Don’t Stress, It’s Under Control: Neural Correlates of Stressor Controllability in Humans"

This supplementary material has been provided by the authors to give readers additional information about their work. Behavioural data and analysis code are available on the Open Science Framework: https://osf.io/8qpme/

**Supplemental Table 1**. Stressor aversiveness ratings – model details

| **Fixed Effects** | | | | | | |
|---|---|---|---|---|---|---|
| *Predictor* | | *Estimate* | *SE* | *95% CI* | *t* | *p* |
| Intercept | | 63.84 | 2.19 | 59.55 – 68.12 | 29.19 | **< .001** |
| Condition | | -0.44 | 0.30 | -1.02 – 0.15 | -1.470 | .143 |
| Run | | -3.71 | 1.09 | -5.84 – -1.58 | -3.414 | **.001** |
| Condition x Run | | -0.22 | 0.30 | -0.80 – -0.37 | -0.725 | .469 |
| **Random Effects** | | | | | | |
| | | *Variance* | | *SD* | *Correlation* | |
| Participant | (Intercept) | 211.18 | | 14.53 | | |
| | Run (Slope) | 49.01 | | 7.00 | 0.42 | |
| **Model Fit** | | | | | | |
| Marginal $R^2$/Conditional $R^2$: 0.046/0.898 | | | | | | |

Model equation: aversiveness ~ condition * run + (1 + run | participant)

**Supplemental Table 2**. Perceived control ratings – model details

| Fixed Effects | | | | | | |
|---|---|---|---|---|---|---|
| *Predictor* | | *Estimate* | *SE* | *95% CI* | *t* | *p* |
| Intercept | | 48.21 | 1.84 | 44.61 – 51.81 | 26.23 | **< .001** |
| Condition | | 16.44 | 1.61 | 13.28 – 19.60 | 10.20 | **< .001** |
| Run | | 0.87 | 1.12 | -1.32 – 3.06 | 0.78 | .442 |
| Condition x Run | | 1.79 | 0.73 | 0.37 – 3.22 | 2.47 | **.014** |
| **Random Effects** | | | | | | |
| | | *Variance* | | *SD* | *Correlation* | |
| Participant | (Intercept) | 128.08 | | 11.32 | | |
| | Condition (Slope) | 93.24 | | 9.66 | -0.06 | |
| | Run (Slope) | 31.99 | | 5.66 | 0.27 -0.04 | |
| **Model Fit** | | | | | | |
| Marginal $R^2$/Conditional $R^2$: 0.385/0.738 | | | | | | |

Model equation: perceived control ~ condition * run + (1 + condition + run | participant)

**Supplemental Table 3**. Stress ratings – model details

| **Fixed Effects** | | | | | | |
|---|---|---|---|---|---|---|
| *Predictor* | | *Estimate* | *SE* | *95% CI* | *t* | *p* |
| Intercept | | 37.89 | 2.45 | 33.08 – 42.70 | 15.45 | **< .001** |
| Condition-CON | | -17.39 | 2.07 | -21.44 – -13.34 | -8.41 | **< .001** |
| Condition-UNCON | | 7.10 | 1.33 | 4.49 – 9.70 | 5.34 | **< .001** |
| **Random Effects** | | | | | | |
| | | *Variance* | | *SD* | *Correlation* | |
| Participant | (Intercept) | 267.24 | | 16.35 | | |
| | Condition-CON (Slope) | 180.09 | | 13.42 | -0.24 | |
| | Condition-UNCON (Slope) | 64.51 | | 8.03 | 0.20 -0.79 | |
| | Run (Slope) | 23.98 | | 4.90 | 0.14 0.15 -0.01 | |
| **Model Fit** | | | | | | |
| Marginal $R^2$/Conditional $R^2$: 0.249/0.855 | | | | | | |

Model equation: stress ~ condition + (1 + condition + run | participant)

**Supplemental Table 4**. Helplessness ratings – model details

| Fixed Effects | | | | | |
|---|---|---|---|---|---|
| *Predictor* | *Estimate* | *SE* | *95% CI* | *t* | *p* |
| Intercept | 45.78 | 2.56 | 40.77 – 50.79 | 17.91 | **< .001** |
| Condition | -10.50 | 1.80 | -14.03 – -6.97 | -5.83 | **< .001** |

**Random Effects**

| | | *Variance* | *SD* | *Correlation* |
|---|---|---|---|---|
| Participant | (Intercept) | 240.03 | 15.50 | |
| | Condition (Slope) | 138.45 | 11.77 | -0.09 |
| | Run (Slope) | 36.61 | 6.05 | -0.03 -0.13 |
| | Stress Duration Difference (Slope) | 88.44 | 9.40 | 0.13 0.51 -0.22 |
| | Condition x Run (Slope) | 8.12 | 2.85 | -0.22 0.46 0.03 |

**Model Fit**

Marginal $R^2$/Conditional $R^2$: 0.167/0.739

Model equation: helplessness ~ condition + (1 + condition * run + stress duration difference | participant)

**Supplemental Table 5**. Reaction times – model details

| | | | | | | |
|---|---|---|---|---|---|---|
| | | **Fixed Effects** | | | | |
| *Predictor* | | *Estimate* | *SE* | *95% CI* | *t* | *p* |
| Intercept | | 769.38 | 6.17 | 757.28 – 781.47 | 124.65 | **< .001** |
| Condition-CON | | 18.65 | 2.55 | 13.65 – 23.65 | 7.31 | **< .001** |
| Condition-UNCON | | -27.04 | 2.09 | -31.13 – -22.94 | -12.94 | **< .001** |
| **Random Effects** | | | | | | |
| | | *Variance* | | *SD* | *Correlation* | |
| Participant | (Intercept) | 1038.02 | | 32.22 | | |
| | Run (Slope) | 84.66 | | 9.20 | 0.37 | |
| | Stress Duration Difference (Slope) | 1502.21 | | 38.76 | -0.02 -0.36 | |
| **Model Fit** | | | | | | |
| Marginal $R^2$/Conditional $R^2$: 0.031/0.116 | | | | | | |

Model equation: reaction time ~ condition + (1 + run + stress duration difference | participant)

**Supplemental Table 6**. Correct responses – model details

| **Fixed Effects** | | | | | | |
|---|---|---|---|---|---|---|
| *Predictor* | | *Estimate* | *SE* | *95% CI* | *z* | *p* |
| Intercept | | 2.05 | 0.11 | 1.84 – 2.26 | 19.12 | **< .001** |
| Condition-CON | | -0.31 | 0.07 | -0.44 – -0.18 | -4.63 | **< .001** |
| Condition-UNCON | | 0.30 | 0.06 | 0.18 – 0.42 | 4.86 | **< .001** |
| **Random Effects** | | | | | | |
| | | *Variance* | | *SD* | *Correlation* | |
| Participant | (Intercept) | 0.17 | | 0.41 | | |
| | Stress Duration Difference (Slope) | 0.36 | | 0.60 | | |
| **Model Fit** | | | | | | |
| Marginal $R^2$/Conditional $R^2$: 0.015/0.062 | | | | | | |
| Model equation: correct ~ condition + (1 + stress duration difference \| \| participant) | | | | | | |

**Supplemental Table 7**. Heart Rate – model details

| | | | | | |
|---|---|---|---|---|---|
| | **Fixed Effects** | | | | |
| *Predictor* | *Estimate* | *SE* | *95% CI* | *t* | *p* |
| Intercept | 64.44 | 1.67 | 61.17 – 67.71 | 38.57 | **< .001** |
| Condition-CON | -0.53 | 0.17 | -0.86 – -0.21 | -3.22 | **.00137** |
| Condition-UNCON | 0.54 | 0.17 | 0.22 – 0.87 | 3.26 | **.00122** |

**Random Effects**

| | | *Variance* | *SD* | *Correlation* |
|---|---|---|---|---|
| Participant | (Intercept) | 105.85 | 10.29 | |
| | Run (Slope) | 8.19 | 2.86 | |
| | Stress Duration Difference (Slope) | 13.72 | 3.70 | |

**Model Fit**

Marginal $R^2$/Conditional $R^2$: 0.002/0.943

Model equation: heart rate ~ condition + (1 + run + stress duration difference | | participant)

**Supplemental Table 8**. Helplessness ratings and ventromedial prefrontal cortex parameter estimates

| **Fixed Effects** | | | | | | |
|---|---|---|---|---|---|---|
| *Predictor* | | *Estimate* | *SE* | *95% CI* | *t* | *p* |
| Intercept | | 46.61 | 2.68 | 41.36 – 51.86 | 17.41 | **< .001** |
| Condition | | -10.00 | 1.10 | -12.12 – -7.89 | -9.28 | **< .001** |
| Beta Weight | | -0.45 | 1.29 | -2.98 – 2.07 | -0.35 | .72535 |
| Condition x Beta Weight | | 3.08 | 1.10 | 0.93 – 5.23 | 2.81 | **.00531** |
| **Random Effects** | | | | | | |
| | | *Variance* | | *SD* | *Correlation* | |
| Participant | (Intercept) | 182.30 | | 13.50 | | |
| | Stress Duration Difference (Slope) | 103.10 | | 10.15 | -0.15 | |
| **Model Fit** | | | | | | |
| Marginal $R^2$/Conditional $R^2$: 0.168/0.442 | | | | | | |

Model equation: helplessness ~ condition * beta weight + (1 + stress duration difference | participant)

**Supplemental Table 9.** Brain activations associated with stress during anticipation

| Region | | MNI coordinates | | | *Z* | $p_{FWE}$ | # voxels |
|---|---|---|---|---|---|---|---|
| **Stress > Baseline** | | | | | | | |
| Rolandic Operandum | L | -50 | -6 | 10 | > 8 | < .001 | 1287 |
| | R | 44 | -4 | 8 | > 8 | < .001 | 916 |
| Hippocampus | R | 30 | -12 | -12 | > 8 | < .001 | 132 |
| | L | -30 | -10 | -12 | > 8 | < .001 | 54 |
| Inferior occipital gyrus | R | 40 | -82 | -6 | > 8 | < .001 | 386 |
| | L | -22 | -86 | -6 | 6.55 | < .001 | 44 |
| Postcentral gyrus | R | 22 | -44 | 74 | 7.40 | < .001 | 176 |
| Paracentral lobule | L | -8 | -18 | 68 | 7.28 | < .001 | 580 |
| Calcarine fissure | R | 14 | -92 | -6 | 6.92 | < .001 | 128 |
| Middle occipital gyrus | L | -24 | -90 | 6 | 6.88 | < .001 | 65 |
| | L | -44 | -86 | 4 | 5.58 | < .001 | 11 |
| Middle cingulate gyrus | L | -6 | -6 | 42 | 6.78 | < .001 | 40 |
| | R | 6 | -6 | 40 | 5.88 | < .001 | 25 |
| Insula | R | 40 | 0 | -6 | 5.91 | < .001 | 27 |
| ventral striatum | | 6 | 0 | -10 | 5.86 | < .001 | 12 |
| temporal pole: superior temporal gyrus | R | 58 | 4 | -8 | 5.83 | < .001 | 44 |
| supplementary motor area | R | 8 | -12 | 80 | 5.75 | < .001 | 12 |

| | | | | | | | |
|---|---|---|---|---|---|---|---|
| Calcarine fissure | L | -12 | -48 | 6 | 5.55 | < .001 | 12 |
| Mediodorsal medial magnocellular thalamus | L | -4 | -20 | 4 | 5.35 | < .001 | 10 |
| **Baseline > Stress** | | | | | | | |
| no suprathreshold clusters ≥ 10 voxels | | | | | | | |

Only maxima of clusters significant at $p_{FWE} < .01$, whole-brain corrected with a cluster-defining threshold of $p = .001$ and at least 10 voxels are reported; smoothness of FWHM = 8.8 x 8.7 x 8.6 mm, volume of 2010.3 resels. L = left; R = right; MNI = Montreal Neurological Institute.

**Supplemental Table 10**. Brain activations related to stressor controllability during anticipation

| Region | | MNI coordinates | | | $Z$ | $p_{FWE}$ | # voxels |
|---|---|---|---|---|---|---|---|
| **CON > UNCON** | | | | | | | |
| Inferior frontal gyrus pars triangularis | R | 54 | 28 | 24 | 6.93 | < .001 | 229 |
| Middle temporal gyrus | R | 58 | -38 | 2 | 6.08 | < .001 | 54 |
| Middle frontal gyrus | R | 42 | 18 | 44 | 6.06 | < .001 | 72 |
| Cerebellum | L | -8 | -80 | -30 | 5.90 | < .001 | 22 |
| Angular gyrus | R | 44 | -58 | 54 | 5.65 | < .001 | 52 |
| **UNCON > CON** | | | | | | | |
| Inferior occipital gyrus | L | -20 | -94 | -6 | > 8 | < .001 | 245 |
| Lingual gyrus | R | 20 | -90 | -6 | > 8 | < .001 | 207 |
| Superior frontal gyrus, medial orbital | L | -6 | 38 | -10 | 6.08 | < .001 | 51 |

Only maxima of clusters significant at $p_{FWE} < .01$, whole-brain corrected with a cluster-defining threshold of $p = .001$ and at least 10 voxels are reported; smoothness of FWHM = 8.8 x 8.7 x 8.6 mm, volume of 2010.3 resels. CON = controllable stress; UNCON = uncontrollable stress; L = left; R = right; MNI = Montreal Neurological Institute.

**Supplemental Table 11.** Local maxima of the cluster in the ventromedial prefrontal cortex (980 voxels) associated with the baseline condition during fixation

| **Region** | | **MNI coordinates** | | | ***Z*** | $p_{FWE}$ |
|---|---|---|---|---|---|---|
| **Baseline > Stress** | | | | | | |
| Ventromedial prefrontal cortex | R | 4 | 62 | -2 | 5.93 | <0.001 |
| | | 2 | 36 | -20 | 5.05 | <0.001 |
| | | 0 | 40 | -20 | 4.97 | <0.01 |
| | | 4 | 48 | -10 | 4.37 | <0.01 |
| | | 0 | 46 | -20 | 4.27 | <0.05 |
| | | 4 | 22 | -18 | 4.23 | <0.05 |
| | L | -6 | 24 | -18 | 5.35 | <0.001 |
| | | -4 | 44 | -8 | 4.24 | <0.05 |
| | | -10 | 44 | -10 | 4.00 | <0.05 |

Only maxima significant at $p_{FWE} < .05$, small-volume corrected with a cluster-defining threshold of $p = .001$ are reported; smoothness of FWHM = 9.0 x 8.9 x 8.8 mm, volume of 33.3 resels. L = left; R = right; MNI = Montreal Neurological Institute.

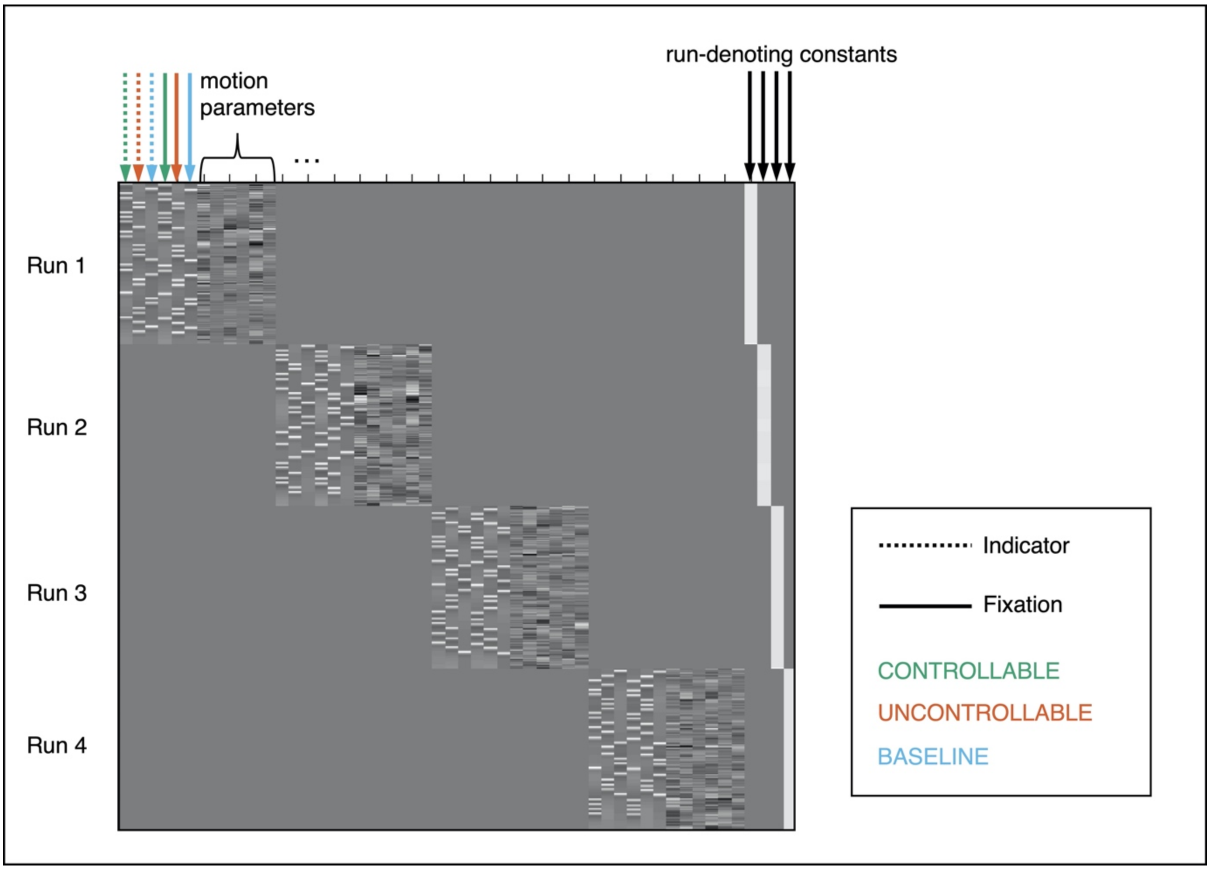


**Supplemental Figure.** First-level model design matrix. All runs comprised the same parameters as visualised for run 1.

**Supplement 13.** Connectivity analysis

We performed a generalised form of psychophysiological interaction analysis (gPPI; McLaren et al., 2012) using Statistical Parametric Mapping (SPM8; The Wellcome Centre for Human Neuroimaging, London, UK; https://www.fil.ion.ucl.ac.uk/spm/software/spm8/). We based the analysis on our general linear model which included six regressors representing the indicator (1500 ms) and fixation phase (4000 ms) of each condition (CON, UNCON, baseline), all convolved with the hemodynamic response function (HRF). However, we focused only on the fixation phase. Physiological regressors were extracted as the 1$^{st}$ eigenvariate of a sphere (6 mm radius) around the first local maximum of the ventromedial prefrontal cortex (vmPFC) cluster which resulted from the CON > UNCON contrast (Figure 5a in main text). Next, PPI regressors were generated by deconvolving the physiological regressors with the HRF, multiplying with each of the condition vectors (fixation only) and reconvolving with the HRF. Thus, six task regressors, three PPI regressors, one physiological regressor, six motion regressors of no interest, and a run-denoting constant comprised the final model. For each participant and run, we then generated the PPI contrasts controllable > uncontrollable stress. The resulting contrast images were entered into a group level analysis and we conducted a one-sample t-test to assess condition-associated modulations of vmPFC-connectivity.
